# Expanding Microgel Parameters to Model the Tumor Microenvironment and Examine Temozolomide Resistance in Glioblastoma

**DOI:** 10.64898/2026.07.08.737105

**Authors:** Brittany A. Payan, Joel Kattoor, Annika Carrillo Diaz de Leon, Gunnar B. Thompson, Tom G. Molley, Kristopher A. Kilian, Jann N. Sarkaria, Brendan A.C. Harley

**Affiliations:** Dept of Bioengineering; Dept of Molecular and Cellular Biology; Dept. Chemical and Biomolecular Engineering; School of Materials Science and Engineering; School of Chemistry; Molecular and Integrative Cystic Fibrosis Research Center University of New South Wales, Australia Sydney, Australia NSW 2052; Dept of Radiation Oncology Mayo Clinic Rochester, MN 55902; Cancer Center at Illinois; Carl R. Woese Institute for Genomic Biology University of Illinois at Urbana-Champaign Urbana, IL 61801

**Keywords:** microgels, glioblastoma, tumor microenvironment

## Abstract

Glioblastoma (GBM) is a highly aggressive brain tumor with a five-year survival rate of less than 5%. The current standard of care established 20 years ago includes maximal surgical resection and administration of alkylating agent temozolomide (TMZ). GBM is highly invasive, and GBM cells that evade surgical resection can become resistant to TMZ and develop new aggressive secondary tumors. Post-relapse there are few treatment options available to patients. Tissue engineering approaches suggest the opportunity to develop in vitro models of the GBM tumor microenvironment that may accelerate the discovery of novel therapies for GBM. Here, we report the adaptation of hydrogel microdroplets (microgels) to encapsulate GBM cells in a tailorable 3D matrix to assess patters of growth and to screen TMZ drug response using patient-derived xenograft (PDX) specimens. We exploit a unique aspect of the microgel system to account for the cellular heterogeneity within the tumor microenvironment (TME). We combine cell-laden microgels generated from TMZ-resistant and TMZ responsive variants of the same PDX specimens to create heterogeneous populations with varying levels of drug sensitivity. We demonstrate a range of drug resistance phenotypes as a function of the ratio of TMZ-responsive to resistance cells and identify the population required for TMZ-resistance to overtake take the response. We then investigate the influence of tumor mimetic shifts in hyaluronic acid bioavailability and hypoxia on patterns of TMZ resistance. We show exposure to matrix-bound hyaluronan increases TMZ resistance and the glioma stem cell population in both cell variants. Lastly, we report an increase in TMZ sensitivity but divergent changes in the GSC subfraction for TMZ resistant vs responsive GBM in the presence of hypoxia. Together, we demonstrate the versatility of cell-laden microgel approach to replicate heterogenous tumor populations, model shifts in the tumor microenvironment, and rapidly screen therapeutic response.

## 1. Introduction

Glioblastoma (GBM) is an aggressive and lethal brain cancer with a five-year survival rate of less than 5%[1]. The standard of care was established in 2005 and includes surgical debulking of the tumor mass, followed by chemotherapy with alkylating agent, temozolomide (TMZ) [2–4]. Although treatment increases median survival, most patients will relapse within 6 months after completing treatment[5]. A particular challenge is the emergence of TMZ-resistant tumors that are more aggressive than the original source and are difficult to treat. A major challenge is the development of appropriate models to investigate therapeutic intervention[6–8] as a function of the emergence of a TMZ-resistant phenotype. Such tools could enable future screening platforms to aid therapeutic discovery efforts for GBM.

The tumor microenvironment (TME) is a complex and dynamic system highly implicated in GBM survival and patient mortality[9]. The TME is a broad term, and its components differ between tissues, but in GBM it is typically characterized by (1) diverse cellular populations and (2) unique chemical signal that amplify an aggressive tumor phenotype. (1) GBM tumors are believed to contain a tumorigenic population, glioma stem cells (GSCs), that show increased levels of self-renewal, proliferation, and invasion linked to poorer patient outcome[10]. These cells are believed to play a major component of TMZ resistance[11–13]. The GBM microenvironment also contains other notable populations such as microglia and macrophages as well as perivascular zones comprised of endothelial cells, astrocytes, and pericytes. (2) GBM exhibits intratumoral hypoxia, a decrease in oxygenation levels, which is often associated with the emergence of a necrotic tumor core[14, 15]. Hypoxia is often implicated in cellular phenotypes tied to GSC maintenance, tumor vascularization, and TMZ resistance[16–18]. And while GBM tumors contain a mixture of fibrillar proteins (e.g., collagens, laminins) and hyaluronic acid (HA), HA in particular has been linked to GBM migration, GSC activity, and chemoresistance[19]. Strategies that offer facile investigation of the coordinated effect of GBM tumor cell heterogeneity, hypoxia, and matrix HA content may help reveal new therapeutic targeting strategies.

Patient-derived xenograft (PDX) models, where patient cancer cells are introduced into immunocompromised mice, have demonstrated the ability to preserve primary tumor characteristics making them an attractive platform to identify novel therapeutics^[20–22]^. PDX models are a predictive and clinically relevant tool. However, the cost and time associated with assays requires large facilities and an established infrastructure. While traditional two-dimensional culture allows rapid screening of therapeutics, it is often limited by the lack of structure and cell-matrix interactions typically seen in vivo. Encapsulation of cells in hydrogels has more recently been used to mimic integral cell-cell and cell-matrix interactions. While these hydrogels have provided insight regarding drug resistance and therapeutic screening^[23–25]^, they most often are conducted using macro-scale constructs (macrogels) that require large amounts of hydrogel material and cells. Further, their large size (1 mm thick; ∼5 mm in diameter) creates diffusional limitations that may impact cellular activity[26, 27]. More recently, cell encapsulation in hydrogel microparticles (microgels) has been implemented as a method to overcome some macrogel limitations. Specifically, microgels can be fabricated rapidly and require lower material demands due to their size (∼10–1000 μm). Their size also aids diffusive transport, even in larger packed beds of microgels. They have successfully been implemented as tumor screening platforms[28] and are capable of identifying novel therapy regimens for patients[29].

We have recently developed a method to encapsulate GBM cells in gelatin maleimide (GelMAL) microgels in which we are able to assess cellular behavior in microgel culture and establish a method to screen therapeutics in a single or metronomic dose regimen[30]. Here, we use this platform to examine clinically relevant patient-derived xenograft (PDX) lines that contain an influential GSC population. We evaluate cellular behavior in microgel culture by characterizing their growth kinetics post-encapsulation and measuring the maintenance of GSC populations. Mixing cell-laden microgels containing TMZ-resistant or TMZ responsive variants of the same PDX specimens[31, 32], we can mimic a spectrum of tumor cellular heterogeneity and identify a critical population fraction of TMZ-resistant cells necessary for persistence. Further, we leverage the microdroplet system to examine the role hypoxia and matrix-bound HA on GBM stem cell fractions and TMZ response. Our findings demonstrate the versatility of cell-laden microgels to rapidly screen therapeutic response and model the heterogeneity of the tumor microenvironment.

## 2. Materials and Methods

### 2.1. Gelatin Maleimide (GelMAL) Synthesis and Characterization

GelMAL was synthesized based on previous methods[33, 34]. Porcine gelatin type A, 300 bloom (Sigma Aldrich, St. Louis, MO) was combined with a 4:5 (v/v) dimethyl sulfoxide (DMSO; Sigma-Aldrich): aqueous medium of pH 6.0 1.0M 2-morpholinoethanesulfonic acid buffer (MES; Gold Biotechnology, St. Louis, MO) at 40 °C until a homogenous mixture was obtained. Next, a 6x excess N-succinimidyl 3-maleimidopropionate (Tokyo Chemical Industry Co., Ltd) was dissolved in DMSO and incorporated into the vial containing the dissolved gelatin. The pH of this reaction was modified to 4.5 using 1.0 N HCl. The reaction was allowed to progress for 24 h at 40 °C and 720 rpm. Then, the solution was dialyzed against distilled water acidified to pH 3.25 using 1.0 N HCl, for 5 days at 40 °C. The resulting product was frozen and lyophilized, then stored at −20 °C. ^1^H NMR was used to assess the degree of functionalization (DOF). Here, GelMAL with a DOF between 45 and 65% was used. The mechanical stiffness of the hydrogels was measured by determining the compressive moduli of GelMAL and HAMAL macrogels via unconfined compression using an Instron 5943 (Instron, Norwood, MA). Samples are compressed at a rate of 0.1 mm/min, then the Young’s modulus is acquired from the linear region of the stress–strain curve (2.5–17.5% strain) utilizing a custom MATLAB (MathWorks, Natick, MA) code.

### 2.2. Flow-focusing Microfluidic Device Construction

Flow-focusing microfluidic devices with a 200 μm nozzle were constructed from polydimethylsiloxane (PDMS) by casting onto SU8 masters using the Sylgard 184 silicone elastomer kit (Dow Corning, Midland, Michigan) [33, 35]. The inlet for gel and cell mixture was incised using a 17 Ga needle, all other ports were incised using a 1 mm biopsy. Post-incision, devices were bonded directly to glass slides after plasma treatment.

### 2.3 Glioblastoma Cell Culture

GBM12 (G12) cell variants (PL5199 and TMZ3080) were provided by Dr. Jann Sarkaria (Mayo Clinic, Rochester, MN) and cultured on laminin-coated (Thermo Fisher Scientific, Waltham, MA) flasks. G12 cells are derived from a male patient and have a classical subtype. They demonstrate epidermal growth factor receptor (EGFR) amplification and O6-methylguanine-DNA methyltransferase (MGMT) is methylated. To create the variants, the TMZ-sensitive cell line, G12, was consistently dosed with TMZ and MGMT became unmethylated, increasing TMZ resistance[31]. This cell line is denoted as G12 TMZ. The G12 cell line was also dosed with a placebo control that remained sensitive to TMZ and is referred to as G12 PL. G12 cells were maintained in StemPro NSC Serum Free Medium (Thermo Fisher Scientific) supplemented with 2% (v/v) GlutaMAX (Thermo Fisher Scientific) and 1% (v/v) penicillin/streptomycin (Thermo Fisher Scientific). This medium formulation is denoted as NSC medium. All G12 cells were cultured in in a humidified 5% CO_2_ incubator at 37 °C, excluding microgels cultured in the hypoxia chamber.

### 2.4 Cell-laden Microgel Fabrication

Microgels were formed using a flow-focusing microfluidic device as previously described[30, 33, 35]. The oil phase was composed of 3 vol% SPAN80 (Sigma-Aldrich) in light mineral oil (Sigma-Aldrich). The crosslinker 1,4-dithiothreitol (DTT) (Sigma-Aldrich) was dissolved at 20 mg/ mL in aqueous solution then emulsified with the oil mixture at a 1:15 (v/v) ratio. 4.8 wt% (w/v) polymer solution was dissolved in PBS with 24% (v/v) OptiPrep (Sigma Aldrich) and 0.12% (w/v) Pluronic F-108 (Sigma Aldrich). G12 cell variants and HCM3 cells were passaged using TrypLE Express (Thermo Fisher Scientific). Cells were washed with PBS three times to remove all remnants of TrypLE and media before keeping on ice until mixed with the polymer precursor.

The polymer precursor and cells were mixed at a 5:1 ratio (final cell concentration: 2 x 10^6^ cells/ mL; final polymer percentage: 4% (w/v)), respectively. The oil phase, DTT crosslinker, and polymer precursor were loaded into syringes then independently injected into the microfluidic device using syringe pumps (Pump 11 Elite, Harvard Apparatus, Holliston, MA) at rates of 0.5 μL/ min (oil mixture), 30 μL/ min (crosslinker emulsion), and 5 μL/ min (polymer precursor + cells), respectively. Microgels were collected for 15 minutes then shaken for 20 minutes at 4 °C to allow completion of crosslinking. Microgels were washed with PBS, 0.1% (v/v) Tween-20 (Fisher Scientific) in PBS, and culture medium. Polymers used to create microgels were GelMAL and hyaluronate maleimide (HAMAL; Creative PEGWorks, Durham, NC). Microgels containing hyaluronic acid were composed of 4% (w/v) GelMAL substituted with 0.05% (w/v) HAMAL. PEG microgel were created with the inclusion of a final concentration of 0.25 mM RGD peptide (MedChem Express, GRGDSPC) to facilitate adhesion to the PEG-4MAL monomer.

### 2.5 Immunofluorescent Staining and Imaging Analysis

Cell-laden hydrogels were fixed in formalin, permeabilized with 0.5% Tween20 (Fisher Scientific), and blocked using 5% donkey serum (Sigma Aldrich) containing 0.1% Tween20. Primary antibodies included glioma stem cell markers Nestin (1:100; Abcam, Cambridge, UK) and SOX2 (1:200; Abcam). Alexa Fluor 594 and 647 were used as secondary antibodies (1:500; Jackson ImmunoResearch, West Grove, PA). Antibodies were conjugated overnight at 4°C. Between antibody incubations hydrogels were washed with 0.1% Tween20 in PBS. Phalloidin conjugated to Alexa Fluor 488 (Thermo Fisher Scientific) was diluted in PBS (1:100) and incubated with the microgels for 2 hours at room temperature. Hoechst 33342 (1:1000; Thermo Fisher) was used as a nuclear marker.

Brightfield images were captured using an EVOS M5000 microscope (Thermo Fisher). Cell-laden microgels were imaged after staining utilizing a confocal laser microscope either Zeiss LSM 710 - Multiphoton Microscope or Zeiss LSM 900 Confocal with Airyscan 2 (Zeiss, Oberkochen Germany). Images were analyzed using ImageJ (NIH, Bethesda, MD). Maximal projection was used to convert the z stack into a 2-dimensional image, then the area of each fluorophore was measure using the ‘Analyze Particles’ function. The overlapping signal was calculated by dividing the marker signal over the actin signal.

### 2.6 Extraction of DNA and RNA

On the respective date of DNA and RNA extraction, GelMAL microgels were dissolved using 50 units/ mL collagenase type IV (Worthington Biochemical Corporation, Lakewood, NJ) on a shaker at 37°C for 20 minutes. The reaction was quenched with PBS + 5% fetal bovine serum (FBS; R&D Systems, Minneapolis, MN). Cell pellets were collected at 300 rcf and stored at - 80°C before extraction. Double-stranded DNA (dsDNA) was extracted using the QIAamp DNA Micro Kit (QIAGEN, Hilden, Germany) following the manufacturer’s protocol. dsDNA was quantified using the Quant-iT PicoGreen dsDNA Assay Kit (Thermo Fisher Scientific), and fluorescence was measured using a Synergy LX Multi-Mode Microplate Reader (BioTek, Winooski, VT) at 485 nm excitation wavelength and 528 nm emission wavelength. RNA was extracted using the RNeasy Micro Kit (QIAGEN) following the manufacture’s protocol.

### 2.7 Gene Expression Analysis

Gene expression of cells cultured in GelMAL microgels and macrogels was evaluated using quantitative reverse transcription polymerase chain reaction (RT-qPCR). RNA samples were reversed transcribed into complimentary DNA (cDNA) using the iScript cDNA Synthesis Kit (Bio-Rad, Hercules, CA) on an Applied Biosystems QuantStudio 7 Flex PCR system (Thermo Fisher, Waltham, MA) with 2× Universal SYBR Green Fast qPCR mix (ABclonal Technology, Woburn, MA) and gene specific primers (Integrated DNA Technologies, Newark, NJ) (Table 1). GAPDH was used as a housekeeping gene. Fold change was calculated using the ΔΔCT method and normalized to expression of two-dimensional (2D) cultured cells.

**Table 1.**
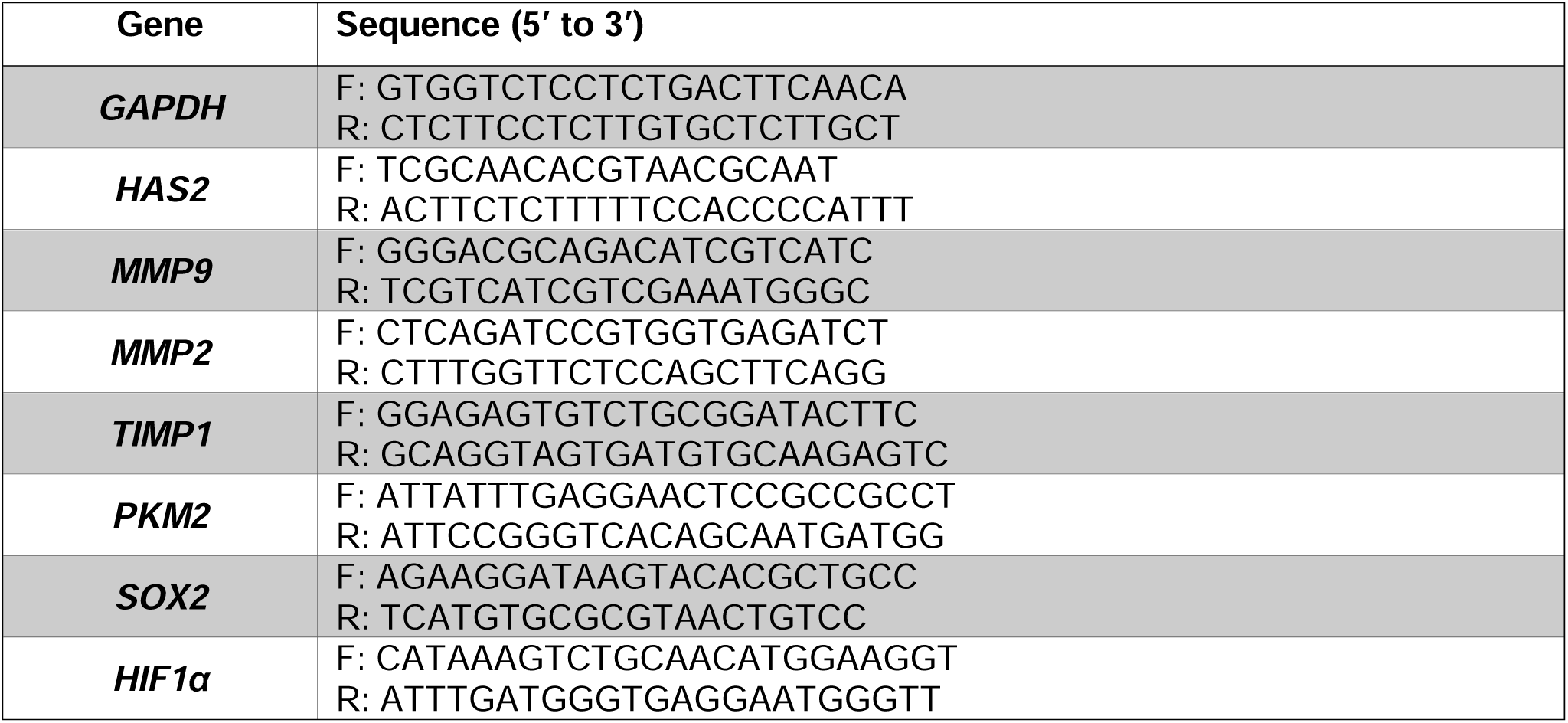
Primer sequences of genes analyzed.

### 2.8 Temozolomide Drug Dosing and IC_50_ Measurement

Cell-laden microgels were cultured in suspension for five days in respective conditions, then manually pipetted into a 384 well plate with each well containing approximately 3 x 10^3^ cells **(Fig S1).** Temozolomide (TMZ, Selleck Chemicals, Houston, TX) dissolved in 100 mM DMSO (Sigma Aldrich) was diluted in culture medium and distributed to the wells. A vehicle control of 0.0001% (v/v) DMSO was used to match the highest volume of DMSO present in the treatment group. Cell-laden microgels were dosed with 0.1, 1.0, 10, 100, and 1,000 μM TMZ. Two days post-treatment, cellular activity was assessed using alamarBlue Cell Viability Reagent (Thermo Fisher Scientific). The reaction was measured by reading the fluorescence using a Synergy LX Multi-Mode Microplate Reader (BioTek) at an excitation wavelength of 530 nm and an emission wavelength of 590 nm. The dose response curve was fit using a non-linear regression model, log(inhibitor) vs response (GraphPad Software Inc., Boston, MA), to measure the TMZ IC_50_ values.

### 2.9 Incorporation of Hyaluronic Acid

Hyaluronic acid (HA) was presented to GBM cells by (1) binding HAMAL into the microgel matrix or (2) adding soluble sodium hyaluronate (41K-65K, Lifecore Biomedical, Chaska, MN) into the culture medium. Matrix bound HA was incorporated by adding 0.05% (w/v) HAMAL into the GelMAL pre-cursor before mixing with G12 cells and fabricating the microgels. To study soluble HA, G12 cells were encapsulated in GelMAL microgels and cultured in medium containing 0.05% (w/v) sodium hyaluronate. GBM cells were exposed to HA throughout the experiment, including post-treatment. All microgels pertaining HA studies were cultured in a humidified 5% CO_2_ incubator at 37 °C.

### 2.10 Integration of Hypoxia

Cell-laden microgels were fabricated under normoxic conditions then exposed to hypoxia. Hypoxia was induced on the microenvironment surrounding GBM-laden microgels by (1) culturing inside a hypoxia chamber (InVivO2; Baker Ruskinn, Sanford, ME) or (2) co-culturing microgels with hypoxic microcapsules. The hypoxic incubator was set to 1% O_2_ and media was equilibrated overnight in the InVivO2 workstation to obtain the desired oxygenation levels.

Hypoxic microcapsules (100 U/ mL) were introduced into the culture media at a 0.5% (v/v) ratio then incubated in a humidified 5% CO_2_ incubator at 37 °C. GBM cells were exposed to hypoxic conditions throughout the experiment, including post-treatment.

### 2.11 Statistics

All statistics were performed using GraphPad Prism. Area, SOX2 and Nestin expression, and metabolic activity of G12 cells are reported as the mean ± standard deviation. The normality of data was verified using the Shapiro–Wilk test, and equality of variance was established using the Brown-Forsythe Test. Samples that passed normality were assessed using the unpaired t-test, one-way analysis of variance (ANOVA) followed by Tukey’s post hoc test, or two-way ANOVA followed by Dunnett’s post hoc test. When data did not satisfy normality, samples were analyzed utilizing a Mann–Whitney test or Kruskal-Wallis test with Dunn’s post hoc test.

Significance is denoted as *p ≤ 0.05, **p ≤ 0.01, ***p ≤ 0.001, ****p ≤ 0.0001. A minimum of three biological samples and three technical replicates were used for all experiments.

## 3. Results

### 3.1 Assessing GBM Cellular Behavior in GelMAL Microgels and Macrogels

We first sought to understand the effect of microgel vs. macrogel culture on GBM cellular behavior. G12 PL GBM patient-derived xenograft (PDX) cells (provided by J. Sarkaria, Mayo Clinic) were encapsulated in maleimide-functionalized gelatin (GelMAL) microgels and macrogels at a concentration (2×10^6^ cells/ mL) to result in an average of 1-3 cells per microgel[30] or 400,000 cells per macrogel (Fig 1A). Actin staining revealed similar G12 PL morphologies in microgel vs. macrogel encapsulation, with cells forming aggregate structures after 7 days of culture (Fig 1B). G12PL cells formed qualitatively larger aggregates in microgels compared to macrogels. We then used RT-qPCR to investigate the differences in cellular function and matrix remodeling between hydrogel models after 7 days in culture. The first panel of genes concentrated on matrix remodeling widely implicated in GBM invasion: hyaluronan synthase 2 (*HAS2*), matrix metallopeptidase 9 (*MMP9*), matrix metallopeptidase 2 (*MMP2*), and tissue inhibitor of metalloproteinases 1 (*TIMP1*). Notably, we observed significant increase in *HAS2* expression (and non-significant increase in *MMP9* expression) from macrogel culture, but significant increase in *MMP2* and decrease in *TIMP1* expression in microgels (Fig 1C). We used a second panel to interrogate genes linked with GBM resistance and survival: pyruvate kinase M2 (*PKM2*), sex-determining region Y (*SRY*)-box transcription factor 2 (*SOX2*), and hypoxia inducible factor 1 alpha (*HIF1a*). We observed significant increases in *PKM2* and *SOX2* in microgel culture, however, an increase in *HIF1a* for cells in larger macrogels (Fig 1D). These results show hydrogel microgels support G12PL culture, matrix remodeling, and heightened expression of genes associated with GBM progression.

**Figure 1.**
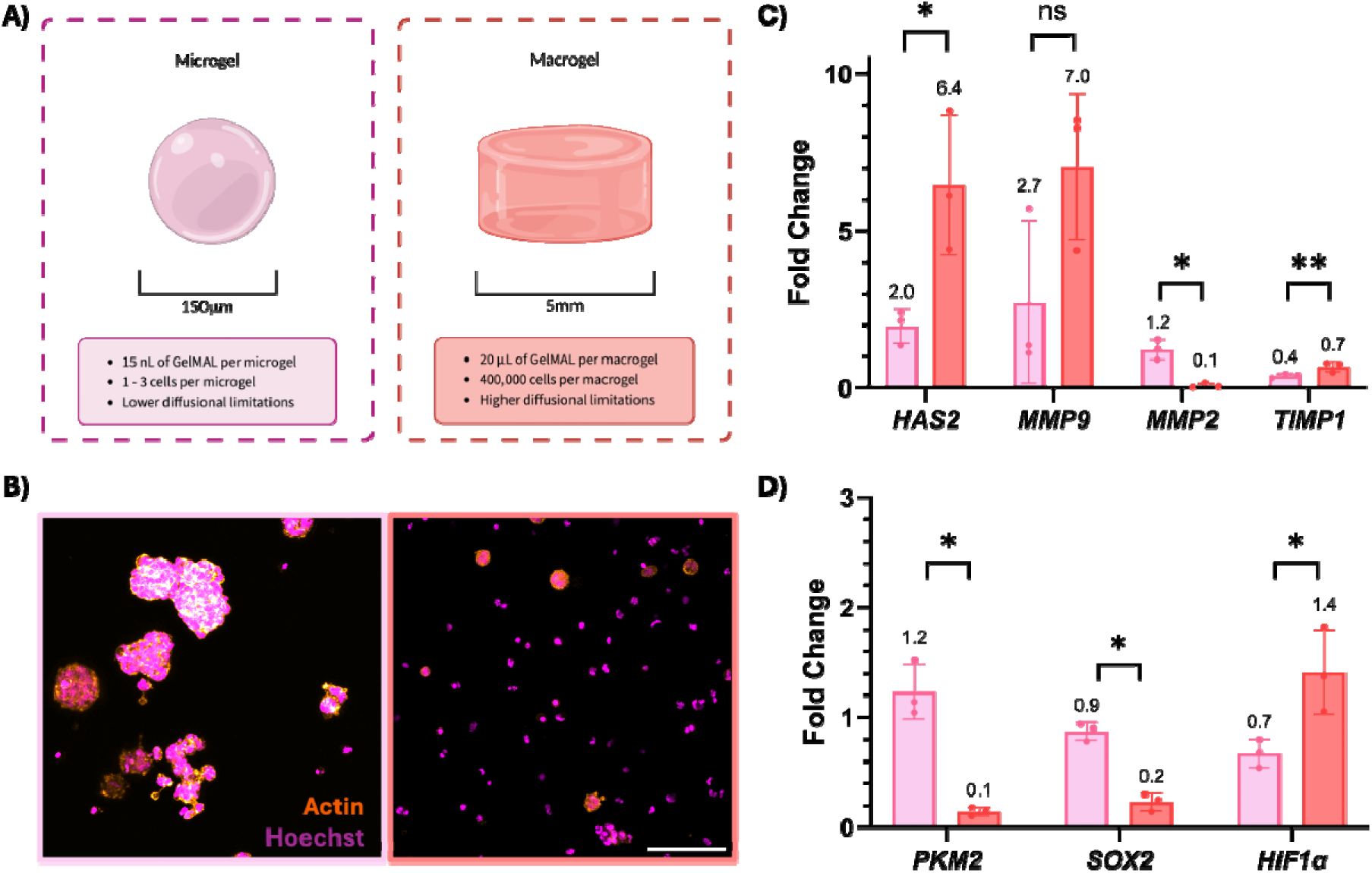
**A)** Schematic comparing the requirements of cell-laden microgel and macrogel models. **B)** Actin stain demonstrated cellular morphology in GelMAL microgel (left) and macrogel (right) culture at day 7. Changes in gene expression for G12 PL cells cultured in GelMAL microgels and macrogels to assesses **C)** matrix remodeling and **D)** cellular function. Scale bar = 100 μm. Bar plot reports mean ± standard deviation. n = 3 independent experiments. Significance is denoted as *p ≤ 0.05, **p ≤ 0.01, ***p ≤ 0.001, ****p ≤ 0.0001.

### 3.2 Microgel Culture of GBM-Responsive and Resistant Patient Derived Xenograft Cells

The G12 PL line is sensitive to temozolomide (TMZ), the main chemotherapeutic utilized in GBM standard of care. However, prior work exposed G12 PL cells to repeated TMZ doses (in vivo) to derive a TMZ-resistant variant, G12 TMZ[31] (Fig S2A). We confirmed that in vitro using 2D culture the G12 PL and G12 TMZ lines display different half maximal inhibitory concentration (IC_50_) values (Fig S2B, C). We subsequently encapsulated G12 PL and G12 TMZ cell lines in GelMAL microgels to define shifts in their growth and stemness. Actin stains demonstrate both G12 variants grow as aggregate structures in GelMAL microgels (Fig 2A, C). Although both lines showed an increase in aggregate size with time, the G12 PL cell aggregates formed significantly larger aggregates by day 5 of culture compared to day 7 by the G12 TMZ cells (Fig 1E). Both aggregates displayed similar levels of expression of GBM stem cell markers *SOX2* and *Nestin* (Fig 2B, D), though *SOX2* rises steadily for the G12 TMZ population after 7-days in culture (Fig 2F). Glioma stem cell marker, *Nestin*, is expressed in a larger fraction of the aggregate populations compared to *SOX2*, and it remains stable over a 7-day culture period for both G12 variants (Fig 2G). These results confirm GBM cells can be cultured in microgels and are able to retain phenotypic markers of GBM stem cells.

**Figure 2.**
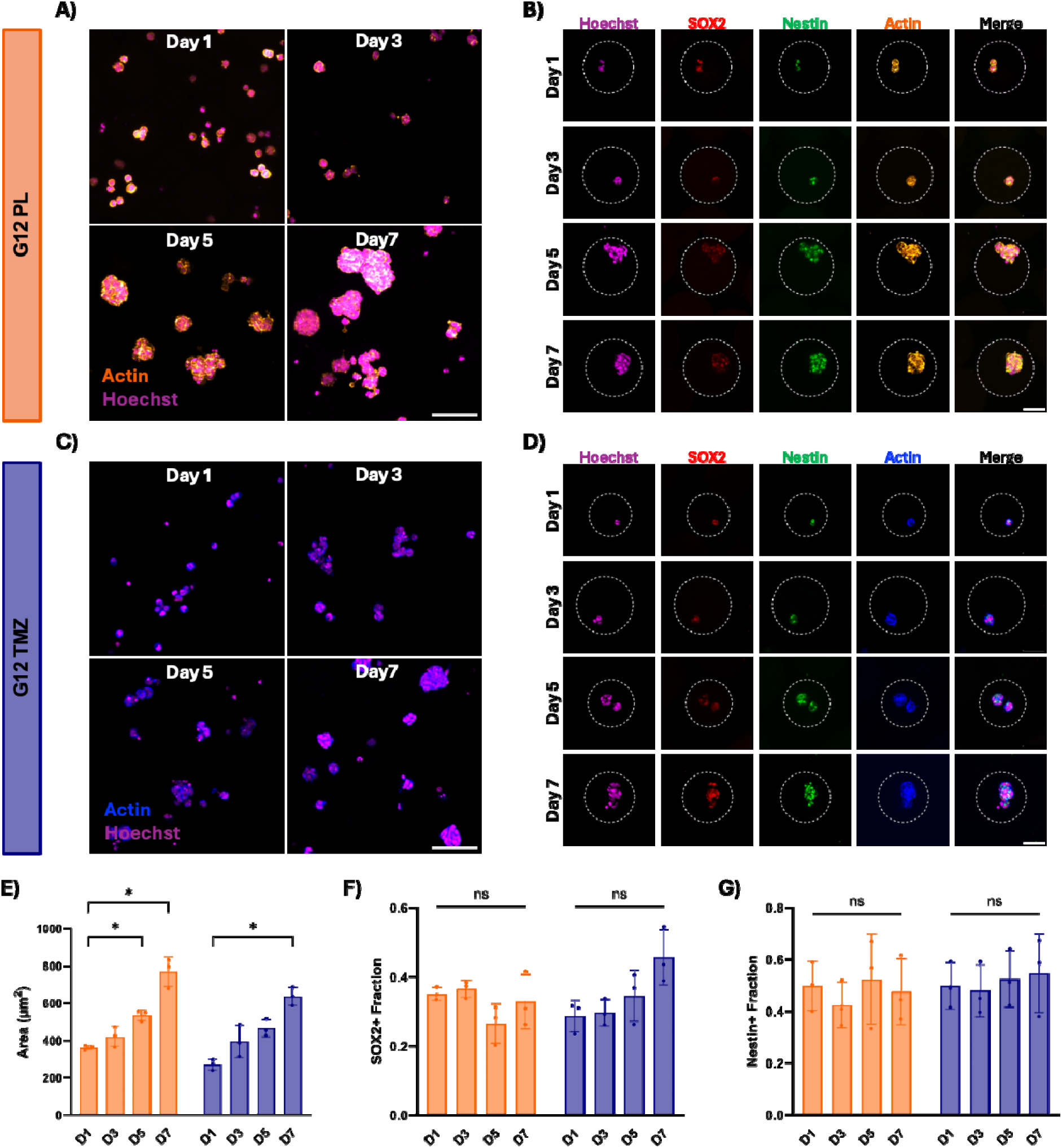
Actin stain demonstrates the size and structure of **A)** G12 PL and **C)** G12 TMZ cells cultured in GelMAL microgels at day 1, 3, 5, and 7. Scale bar = 100 μm. Immunofluorescent staining of markers SOX2 and Nestin in **B)** G12 PL and **D)** G12 TMZ cells in GelMAL microgels at day 7. Scale bar = 50 μm. **E)** Average surface area of G12 cellular aggregates in GelMAL microgels at day 1, 3, 5, and 7. Quantification of cellular populations expressing **F)** SOX2 and **G)** Nestin in GelMAL microgel culture at day 1, 3, 5, and 7. Bar plot reports mean ± standard deviation. n = 3 independent experiments. Significance is denoted as *p ≤ 0.05, **p ≤ 0.01, ***p ≤ 0.001, ****p ≤ 0.0001.

### 3.3 Modeling GBM Tumor Heterogeneity Utilizing Cell-Laden Microgels

Next, we sought to establish a model of heterogeneous tumor phenotypes using mixed populations of G12-laden microgels. We allowed G12 PL and G12 TMZ-laden microgels to culture independently for 5 days, before mixing the variants to build heterogenous populations (100:0, 75:25, 50:50, 25:75, 0:100; G12PL:G12TMZ). Mosaic microgel cultures were then administered a single dose of TMZ with response recorded 2-days post-treatment (Fig 3A). All mixtures retained high metabolic activity at a 10 μM TMZ dose, suggesting cells were not responsive to a dose at or below physiologically relevant levels (Fig 3B). At a supraphysiological dose, mosaic mixtures of TMZ-resistant and TMZ-responsive cells reveal the emergence of unique patterns of TMZ resistance (Fig 3C). Examining the effect of TMZ dose on mosaic cultures, we observed lower sensitivity to increasing TMZ doses (10, 100, 1000 μM) on metabolic activity in mosaic cultures containing higher fractions of TMZ responsive microgels (Fig 3E). As the fraction of TMZ-resistant PDX cells increases (25:75, 0:100; G12PL:G12TMZ), the mosaic cultures exhibit no significant effect of increasing TMZ dosages on the composite metabolic activity of the culture (Fig 3E). Interestingly, the mosaic cultures each demonstrate unique IC_50_ curves (Fig 3F) that reveal higher IC_50_ values with increasing levels of G12 TMZ microgels (277.3 to 1100.9 μM; Fig 3D). The loss of TMZ sensitivity occurs at the transition of the G12 TMZ fraction 50 to 75% (increase IC_50_ value from 492.2 to 801.7 μM) of the overall culture. While this data reveals a facile way to rapidly generate a range of mosaic GBM cultures with control over the relative fraction of TMZ resistant cells that display the emergence of complex composite TMZ resistant phenotypes, we chose a single mixed culture (50:50 G12PL:G12TMZ) to explore shifts in GMB cell phenotype in response to other key parameters inspired by the GBM tumor microenvironment.

**Figure 3.**
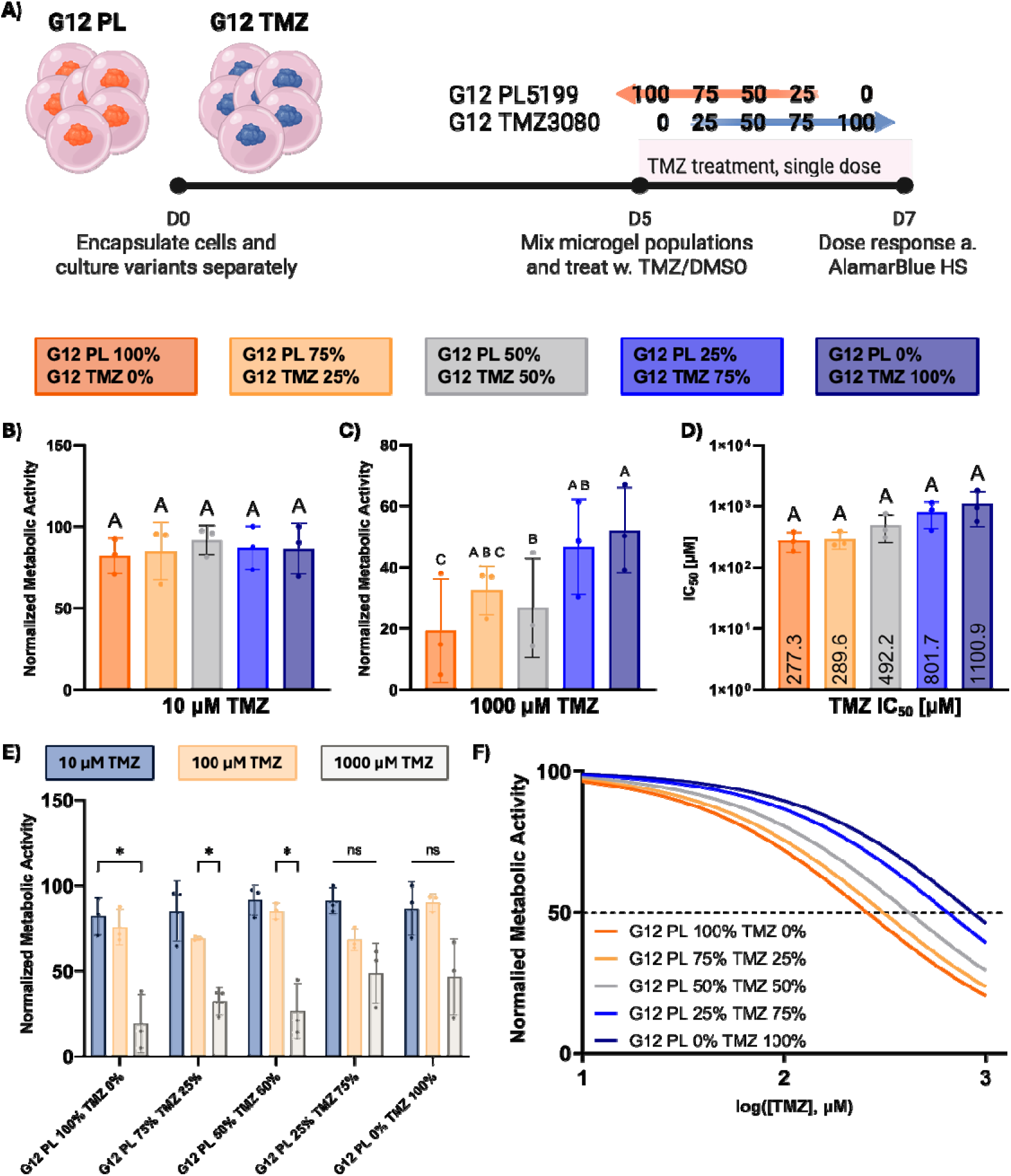
**A)** Methods schematic, G12 PL and G12 TMZ cells are encapsulated in GelMAL microgels and cultured independently for 5 days. On day 5, cell-laden microgels of both populations are mixed at varying ratios (PL:TMZ; 100:0, 75:25, 50:50, 25:75, and 0:100) to model tumor heterogenous populations. After mixing, cells are treated with a single dose of TMZ [0-1,000 μM]. Changes in cellular behavior are assessed 2-days post-treatment via alamarBlue. Treatment response from heterogenous populations exposed to **B)** 10 μM and **D)** 1,000 μM TMZ. **E)** Compilation of cellular response from heterogeneous populations to 10, 100, and 1,000 μM TMZ. **D)** IC_50_ values reported for the 100:0, 75:25, 50:50, 25:75, and 0:100 heterogenous populations. **F)** TMZ IC_50_ curves are derived from treatment response of heterogeneous populations. Bar plot reports mean ± standard deviation. n = 3 independent experiments. Significance is denoted as *p ≤ 0.05, **p ≤ 0.01, ***p ≤ 0.001, ****p ≤ 0.0001.

### 3.4 Incorporating Hyaluronic Acid in Microgel Culture Increases TMZ Resistance in GBM

We subsequently explored shifts in GBM cohort activity in response to matrix-bound hyaluronic acid (HA). We characterized G12 cellular growth and stemness in microgel culture for up to 7 days in either conventional 4 wt% (w/v) GelMAL microgels or HAMAL microgels formed as a mixture of 0.05% (w/v) maleimide-functionalized HA with 3.95 wt% (w/v) GelMAL. Actin staining reveals G12 PDX cells continue to form aggregates with similar morphologies in HAMAL microgels (Fig 4A). Inclusion of HA significantly decreases the size of aggregates formed for G12 variants (Fig 4B). We did not observe differences in the expression of the GBM stem cell marker *SOX2* in GelMAL vs. HAMAL microgels for either G12PL or G12TMZ variants (Fig 4C). However, we observed a significant increase in the expression of the GBM stem cell marker *Nestin* in HAMAL microgels (vs. conventional GelMAL microgels) for both G12 PL and G12 TMZ variants (Fig 4D).

**Figure 4.**
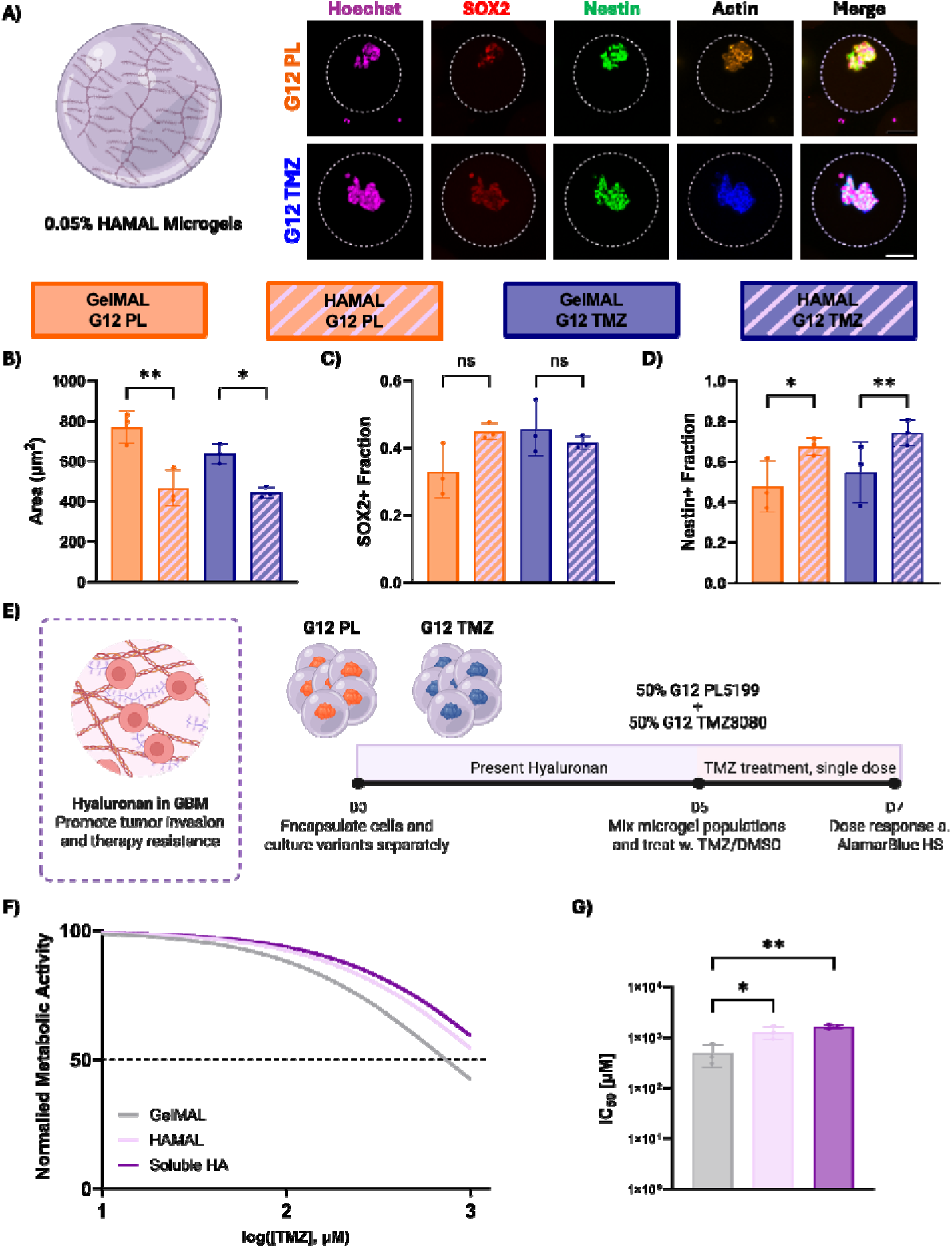
**A)** Actin, SOX2, and Nestin stain of G12 PL (top row) and G12 TMZ (bottom row) cells on day 7 in GelMAL microgels with 0.05% HAMAL incorporated into the matrix. **B)** Average surface area of G12 cellular aggregates in GelMAL microgels with or without HAMAL at day 7. Quantification of cellular populations expressing **C)** SOX2 and **D)** Nestin in GelMAL microgel with or without HAMAL at day 7. **E)** Methods schematic, G12 PL and G12 TMZ cells are encapsulated in GelMAL microgels with matrix bound 0.05% HAMAL or 0.05% soluble HA substituted into the media for 5 days. On day 5, cell-laden microgels of both populations are mixed at a PL:TMZ; 50:50 ratio. After mixing, cells are treated with a single dose of TMZ [0-1,000 μM]. Changes in cellular behavior are assessed 2-days post-treatment via alamarBlue. **F)** TMZ IC_50_ curves are derived from treatment response of 50:50 populations in the presence of HA. **G)** IC_50_ values reported for the 50:50 population in GelMAL, HAMAL, or GelMAL microgels with soluble HA. Scale bar = 50 μm. Bar plot reports mean ± standard deviation. n = 3 independent experiments. Significance is denoted as *p ≤ 0.05, **p ≤ 0.01, ***p ≤ 0.001, ****p ≤ 0.0001.

We then characterized the effect of HA on TMZ resistance. We independently cultured G12 PL and TMZ variants in conventional GelMAL microgels, in HAMAL microgels, or in GelMAL microgels in the presence of 0.05% (w/v) soluble HA for 5 days before combining microgel populations to form the 50:50 G12PL:G12TMZ cultures that were then treated with a single dose of TMZ (Fig 4E). After two days of culture, we used changes in cohort metabolic activity to calculate IC_50_ curves that revealed an increase in TMZ resistance as a result of GBM exposure to soluble or matrix-immobilized HA (Fig 4F). The TMZ IC_50_ value (50:50 G12PL:G12TMZ microgel culture) increased significantly from 492.2 to 1292 μM in HAMAL microgels and 1644 μM in GelMAL microgels in the presence of soluble HA (Fig 4G).

### 3.5 Integrating Hypoxia in Microgel Culture Decreases TMZ Resistance in GBM

Hypoxia is a known feature of the GBM TME that can affect GBM cell activity and drug response. Tissue engineering approaches to investigate the role of hypoxia typically rely on culture in dedicated hypoxia incubators. We first cultured G12-laden (G12 PL or G12 TMZ) microgels in a hypoxic incubator at a consistent oxygen level of 1%, a significant reduction compared to traditional incubators (15-21% oxygen levels[36]). We characterized G12 growth and stemness expression after 7 days in culture. Actin staining revealed both G12 variants grow as (smaller) aggregates and retain the expression of glioma stem markers, *SOX2* and *Nestin* (Fig 5A). While hypoxia did not significantly reduce the size of G12 PL aggregates, G12 TMZ aggregates were significantly smaller in GelMAL microgels exposed to hypoxia (Fig 5B). We observed more complex shifts in GBM stem cell phenotypes, with G12 PL cells cultured under hypoxia, showing significant increases in expression of *SOX2* and *Nestin* (Fig 5C, D), while G12 TMZ cells exhibited a significant decrease in *SOX2* expression under hypoxia (Fig 5C, D).

**Figure 5.**
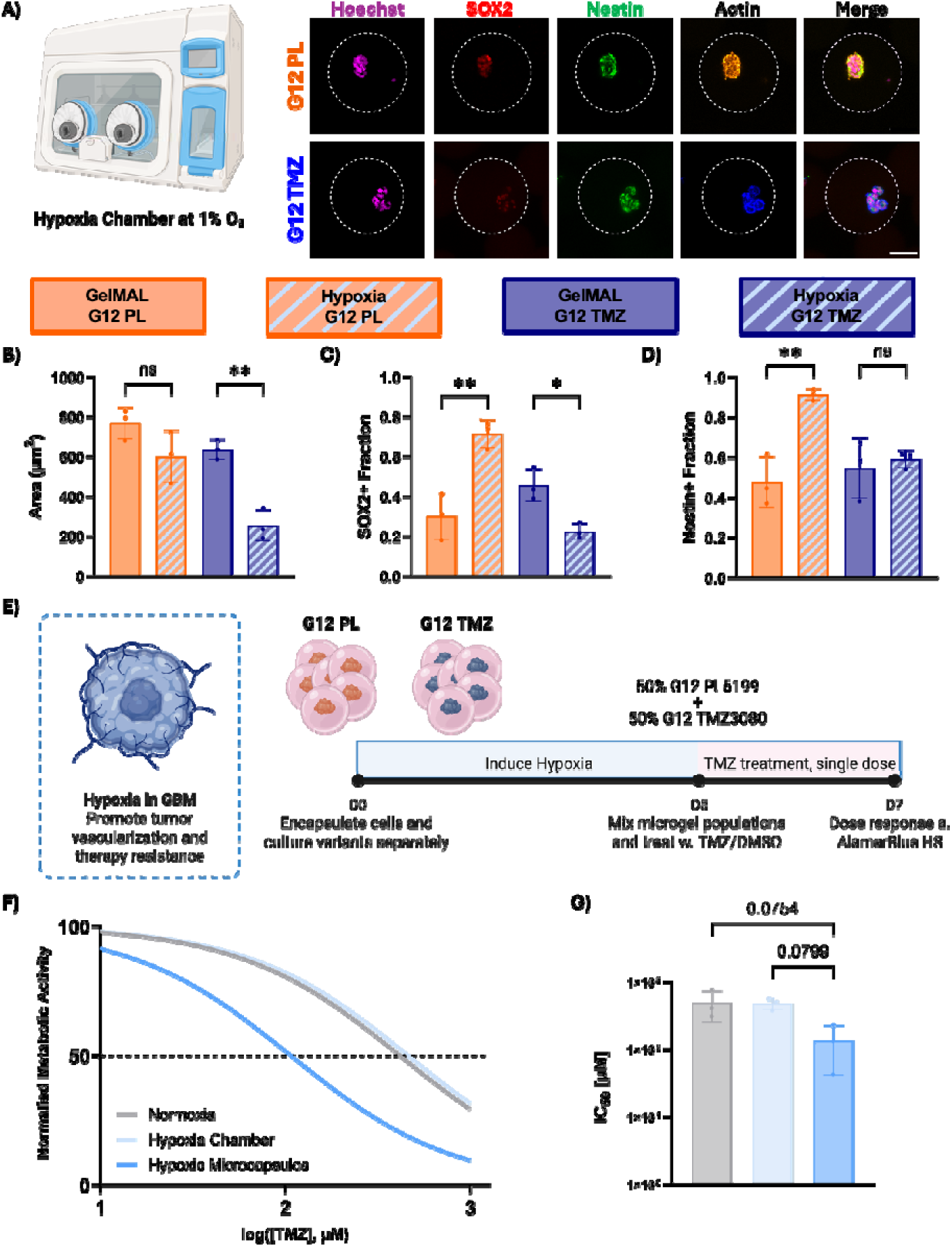
**A)** Actin, SOX2, and Nestin stain of G12 PL (top row) and G12 TMZ (bottom row) cells on day 7 in GelMAL microgels cultured in a hypoxic chamber (1% O_2_). **B)** Average surface area of G12 cellular aggregates in GelMAL microgels cultured in a traditional incubator or a hypoxia chamber at day 7. Quantification of cellular populations expressing **C)** SOX2 and **D)** Nestin in GelMAL microgel cultured in a traditional incubator or a hypoxia chamber at day 7. **E)** Methods schematic, G12 PL and G12 TMZ cells are encapsulated in GelMAL microgels and cultured in a hypoxia chamber (1% O_2_) or co-cultured with 0.5% hypoxic beads in a traditional incubator for 5 days. On day 5, cell-laden microgels of both populations are mixed at a PL:TMZ; 50:50 ratio. After mixing, cells are treated with a single dose of TMZ [0-1,000 μM]. Changes in cellular behavior are assessed 2-days post-treatment via alamarBlue. **F)** TMZ IC_50_ curves are derived from treatment response of 50:50 populations in the presence of hypoxia. **G)** IC_50_ values reported for the 50:50 population cultured in GelMAL microgels under normoxic condition, or hypoxic conditions in a hypoxia chamber or with hypoxic microcapsules in a conventional incubator. Scale bar = 50 μm. Bar plot reports mean ± standard deviation. n = 3 independent experiments. Significance is denoted as *p ≤ 0.05, **p ≤ 0.01, ***p ≤ 0.001, ****p ≤ 0.0001.

We then defined the effect of hypoxia on TMZ resistance for 50:50 G12PL:G12TMZ cohorts. We incubated the G12PL and G12TMZ variants independently in a hypoxia chamber or in a regular incubator. After 5 days in culture, G12 variants were mixed at a 50:50 ratio and treated with a single dose of TMZ then maintained in culture for two days (Fig 5E). Despite the changes in cellular behavior in the presence of hypoxia, IC_50_ curves reveal a similar response from both normoxic and hypoxic (1% O_2_) samples (a slight decrease from 492.2 to 486.2 μM).

Manipulation of cultures in a conventional hypoxia chamber is difficult and requires significant specialized equipment. As a result, we explored an alternative path to create hypoxic cultures that is well suited to the granular nature of the mosaic cultures formed from mixtures of GBM-laden hydrogel microdroplets. G12-laden microgels were cultured in media containing 0.5% (v/v) gelatin-based hypoxic microcapsules (Hypoxycaps; d_avg_ = 60 μm[37]) designed to enzymatically consume oxygen in order to create a local hypoxic environment in conventional incubator culture (Fig S4A). After 5 days in normoxia or hypoxia preconditioning, we generated 50:50 G12PL:G12TMZ microgel cultures entirely maintained in a conventional incubator. The oxygen concentration of media samples from GelMAL-hypoxic microcapsules cultures ranged from 1.746 to 3.233 O_2_ mg/ L (Fig S4B). Notably, the microcapsules exacerbated the hypoxic preconditioning phenotype and we observed a significant decrease in TMZ IC_50_ value for GelMAL-hypoxic microcapsules (135.3 μM) versus conventional microgel culture (492.2 μM).

## 4. Discussion

GBM accounts for a large percentage of primary brain tumors, and the prognosis has remained unchanged for the past two decades[9]. This is in part due to the slow progress of therapeutics from lab to clinic. Currently, there is a critical need for a platform that can recapitulate the local microenvironment and reliably evaluate compounds in a timely manner. PDX models are costly and require large infrastructure, limiting their use for screening compound libraries in a high-throughput method^[38]^. Hydrogels are able to recapitulate essential cell-cell and cell-matrix interactions by enveloping cells in a highly tunable matrix in an accessible manner[39].

Conventional bulk hydrogels, however, possess scalability limitations as they require large amounts of hydrogel material and cellular population demands. Cell-laden microgels suggest an avenue to overcome previous limitations and allow rapid screening of therapeutics in a relevant matrix[29, 30]. Being able to evaluate the effect of specific factors of the TME in a rapid drug screening platform will enhance drug discovery methods and aid in the development of targeted therapeutics for GBM.

We first sought to understand essential differences in cellular behavior of GBM cells encapsulated in GelMAL microgels and macrogels. We encapsulated PDX cell line, GL12 PL, in the respective hydrogel model and assessed cellular behavior after culturing for 7 days.

Morphology reveals the formation of similar cellular aggregates in both models; however, structures are larger in microgel culture suggesting an enhance proliferative profile in microgel culture. This may be of benefit for cellular expansion of low volume patient tumors, where biopsy samples contain a low quantity of cancer cells[40]. Evaluation of *MMP9*, showed similar levels in both models suggesting GBM cells express a basal level of degradation of the matrix that is unaffected by the hydrogel size. However, an increase in *MMP2* and decrease of *TIMP1*, markers with an inverse relationship, corroborate cells in microgels possess significantly higher degradation ability. The upregulation of *HAS2* in macrogels compared to microgels, suggests GBM cells are focusing on hyaluronan synthase-mediated invasion over degradation of the matrix in macrogel culture. Overall, these results demonstrate essential differences in GBM behavior due to the hydrogel size that should be considered and chosen based on the objective of the study.

Having understood the fundamental advantages of microgel culture, we encapsulated PDX cell line variants, G12 PL and G12 TMZ, and characterized their growth in GelMAL microgels. Utilizing PDX cell lines allows us to study primary characteristics of the original tumor as well as assess for patient variability. Both variants continually increased in size over the span of 7 days, retaining viability and proliferative capacity post-encapsulation. We also evaluated GSC marker expression of *SOX2* and *Nestin* to assess the compatibility of our platform in the maintenance of a highly sensitive population. *SOX2* is known to promote dedifferentiation and plays a critical role in chemoresistance[41, 42]. *Nestin* is a regulator of growth in GBM and plays a large role in invasion and migration[43, 44]. Both G12 lines encapsulated in GelMAL microgels consistently expressed *SOX2* and Nestin throughout the 7 days of culture, demonstrating our ability to maintain sensitive populations in microgel culture.

Granular hydrogels possess various advantages, including the temporal control for creating mosaic cultures. We wished to exploit these properties to simulate heterogenous tumor and better understand TMZ resistance. The encapsulation of separate cellular populations in microgels allowed us to easily mix cell-laden microgels and create diverse tumor populations. To create a baseline response of these synthetic tumor populations, we simplified our approach and cultured each population independently (G12 PL and G12 TMZ) from one another until the day of treatment with TMZ. We were then able to record a rapid response from the cells without the competition or crosstalk between populations before treatment. We created G12PL:G12TMZ ratios, 100:0, 75:25, 50:50, 25:75, and 0:100 to assess the contribution of individual populations to TMZ resistance. As expected, increasing the G12 TMZ population decreased the effect of TMZ even at large supraphysiological doses [100 μM and 1,000 μM TMZ]. The derivation of TMZ IC_50_ after 2 days of exposure, demonstrated a steep increase in TMZ resistance with higher G12 TMZ population fractions. The largest jump in TMZ resistance is seen as the G12 populations PL:TMZ are increased from 50:50 to 25:75. Therefore, we chose the 50:50 population to further explore the effect of the TME on TMZ resistance.

Hyaluronic acid (HA, interchangeably referred to as hyaluronan) is a principal component of the native brain ECM, and has been noted to accumulate abnormally in GBM tumors[45]. It is a crucial factor for GBM invasion[46] and is highly correlated with therapeutic resistance[47]. We decided to incorporate HAMAL, a matrix bound version of HA into the microgel composition to assess changes in GBM behavior. Inclusion of HAMAL significantly decreased the size of G12 aggregates, indicating a transition to a migratory phenotype over a proliferative state[48]. This was further supported by the upregulation of *Nestin* in both cell lines, a protein that plays a large role in migration and has previously been reported to increase in the presence of HA[49]. Lastly, we assessed the impact of HA in TMZ resistance by culturing G12-laden microgels in the presence of matrix bound HA (HAMAL) or solubilized HA substituted into the culture medium.

As expected, the TMZ IC_50_ value for the 50:50 populations increased, showing a 3-4-fold increase in the presence of HA. Interestingly, the way HA was presented to the cells did not show differences in TMZ response, demonstrating either method is suitable to screen compounds in the presence of HA.

Lastly, we explored the role of hypoxia another factor of the TME implicated in GBM progression. Typically hypoxia develops as tumors grow in size and deplete the nutrients and oxygen required for tumor growth[50]. Hypoxia has been correlated to GSC maintenance[17, 51], making it an important factor to study in the development of therapeutics targeting GSC survival. We cultured the G12 variants in GelMAL microgels inside a hypoxia chamber (1% O_2_) to mimic the low oxygenation cells may be exposed to in the brain[52]. Both G12 PL and G12 TMZ cells formed aggregate structures in the presence of hypoxia, however, G12 TMZ cells show a significant decrease in size demonstrating lower proliferative capabilities. TMZ-sensitive G12 PL cell line behaved according to the classic phenotype established in previous literature, demonstrating an increase in GSC populations under hypoxic conditions. On the other hand, G12 TMZ cellular behavior diverged and displayed significantly low expression of SOX2. We then assessed the impact of hypoxia on TMZ resistance by pre-conditioning the cells by culturing them under hypoxic conditions then dosing with TMZ. The 50:50 samples cultured at 1% O_2_ in a hypoxia chamber demonstrated a slight decrease in IC_50_ compared to the normoxic conditions. To assess hypoxia in a more practical manner, we utilized hypoxic microcapsules to induce a hypoxic environment in a conventional incubator. The IC_50_ derived from 50:50 samples co-cultured with hypoxic microcapsules, showed a significantly lower value compared to the normoxic conditions. This is likely due to the lower oxygenation achieved from the microcapsules, as well as the depletion of glucose in the process[37]. The decrease in IC_50_ values aligns closely with the depletion of resistant G12 TMZ populations, demonstrating their inability to contribute to the resistance reading during hypoxia. Further studies need to be completed to determine the G12 TMZ cellular state in hypoxic conditions. Overall, this study has emphasized the importance of the TME in TMZ resistance and has provided a method to account for these variables in an in vitro drug screening platform. These results support the future of cell-laden microgels as a platform for drug discovery as well as a method to study specific factors of the TME.

## 5. Conclusions

Hydrogel macrogels have proven to be a useful tool to assess GBM invasion, migration, and therapeutic response. However, due to their large size they possess diffusional limitations and unconventional material requirements. Cell-laden microgels have been utilized as an alternative to encapsulate cells in a tailorable matrix in a scalable manner. Previously, we demonstrated the ability to rapidly assess drug-response and model different dosing regimens using GBM cell-laden microgels. Here, we expand upon this work and demonstrate the utility of this platform to study factors of the TME on therapy response. We assess patient compatibility by encapsulating PDX cell lines and demonstrating the maintenance of GSC populations over 7 days. We model tumor heterogeneity by mixing cell-laden microgels of TMZ sensitive and TMZ resistant populations. We identify the 50:50 TMZ sensitive:resistant population as the requirement to shift towards a TMZ resistant phenotype. We assess changes to TMZ response in this 50:50 population by integrating factors of the TME. We demonstrated that hyaluronan availability increases TMZ resistance and hypoxia increases TMZ sensitivity in microgel culture. This work demonstrates the versatility of microgel culture and the advantages of utilizing cell-laden droplets as a drug screening platform.

## Acknowledgements

We acknowledge the following institutes for access to their facilities and services: the Roy J. Carver Biotechnology Center at UIUC, the Tumor Engineering and Phenotyping Core at the Cancer Center at Illinois, and the Carle R Woese Institute for Genomic Biology. Research reported in this publication was supported by the National Cancer Institute of the National Institutes of Health under Award Number R01 CA256481 (BACH), the National Institute of Diabetes and Digestive and Kidney Diseases of the National Institutes of Health under Award Number 2 R01 DK099528 (BACH), as well as the National Institute of Biomedical Imaging and Bioengineering of the National Institutes of Health under Award Number T32EB019944. This manuscript is the result of funding in whole or in part by the National Institutes of Health (NIH). It is subject to the NIH Public Access Policy. Through acceptance of this federal funding, NIH has been given a right to make this manuscript publicly available in PubMed Central upon the Official Date of Publication, as defined by NIH. The content is solely the responsibility of the authors and does not necessarily represent the official views of the NIH. The authors are also grateful for additional funding provided by the Department of Chemical & Biomolecular Engineering, the Carl R. Woese Institute for Genomic Biology, the Cancer Center at Illinois, and the Illinois Scholars Undergraduate Research Program at the University of Illinois Urbana-Champaign.

## Contributions (CRediT: Contributor Roles Taxonomy [53, 54])

**B.A. Payan:** Conceptualization, Data Curation, Formal Analysis, Visualization, Investigation, Methodology, Writing – original draft, Writing – review & editing.

**J. Kattoor:** Data Curation, Investigation

**A. Carrillo Diaz De Leon:** Data Curation, Investigation

**G.B. Thompson:** Methodology

**T.G. Molley:** Methodology

**K.A. Kilian:** Resources

**J.N. Sarkaria:** Resources

**B.A.C. Harley:** Conceptualization, Resources, Project administration, Funding acquisition, Supervision, Writing – review & editing.

