## Supplemental Info for "Expanding Microgel Parameters to Model the Tumor Microenvironment and Examine Temozolomide Resistance in Glioblastoma"

^1^ Dept of Bioengineering

^2^ Dept of Molecular and Cellular Biology

^3^ Dept. Chemical and Biomolecular Engineering

^8^ Cancer Center at Illinois

^9^ Carl R. Woese Institute for Genomic Biology

University of Illinois at Urbana-Champaign

Urbana, IL 61801

^4^ School of Materials Science and Engineering

^5^ School of Chemistry

^6^ Molecular and Integrative Cystic Fibrosis Research Center

University of New South Wales, Australia

Sydney, Australia NSW 2052

^7^ Dept of Radiation Oncology

Mayo Clinic

Rochester, MN 55902

**Corresponding Author:**

B.A.C. Harley

Dept. of Chemical and Biomolecular Engineering

Cancer Center at Illinois

Center for Gender & Sex in Health

Carl R. Woese Institute for Genomic Biology

University of Illinois at Urbana-Champaign

110 Roger Adams Laboratory

600 S. Mathews Ave.

Urbana, IL 61801

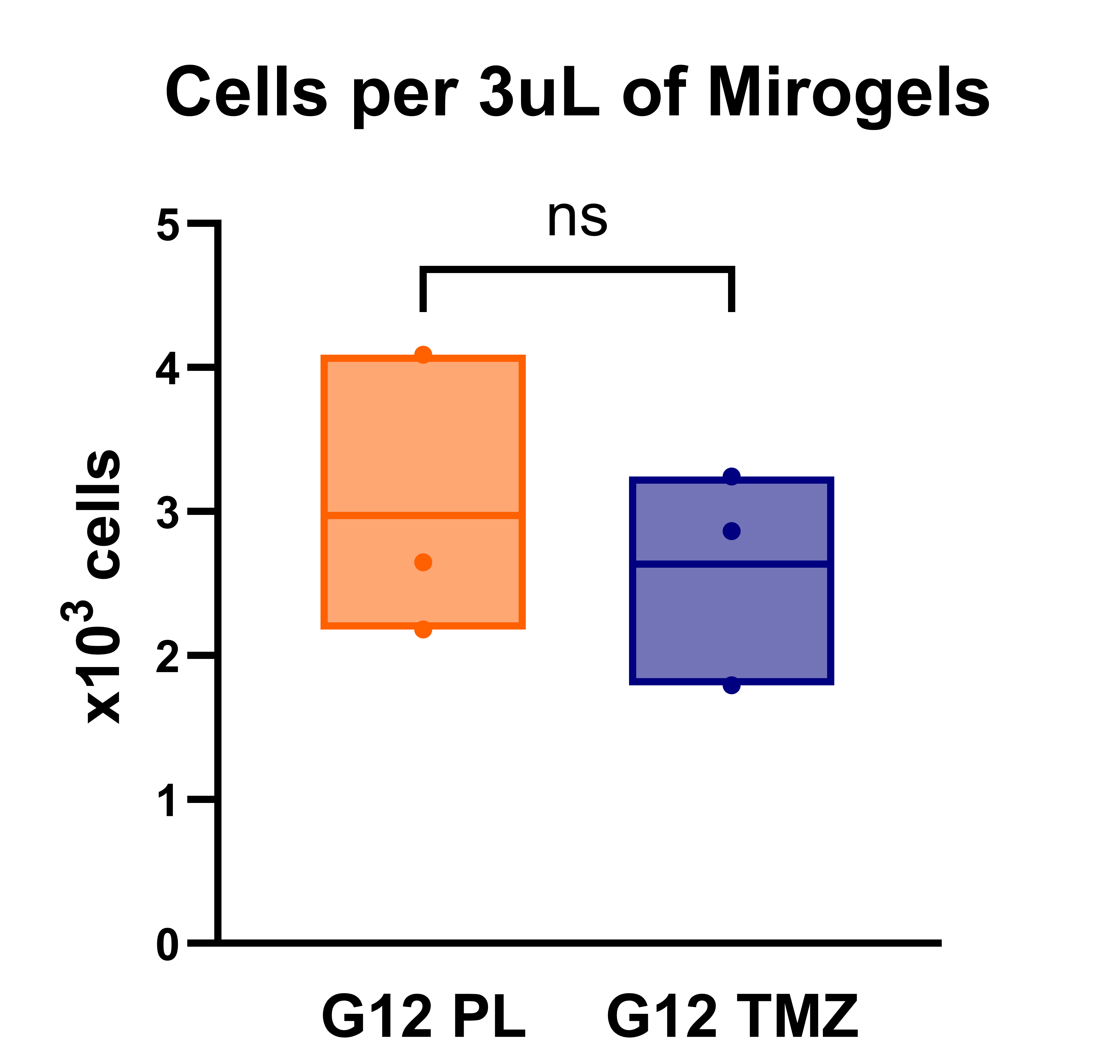

**Figure S1.** Quantity of cells in a 3 μL sample of G12-laden microgels. G12 PL microgels contain approximately 2.973 x 10^3^ cells per sample. G12 TMZ microgels contain approximately 2.634 x 10^3^ cells per sample.

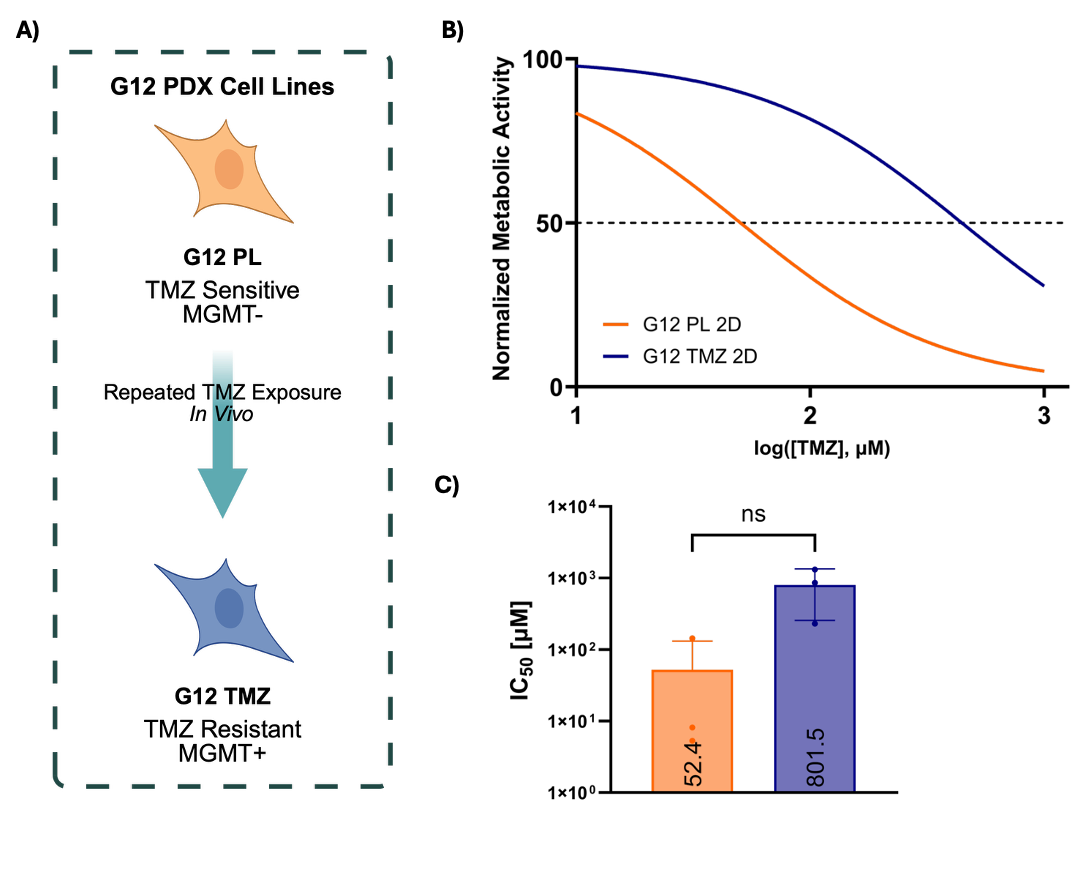

**Figure S2. A)** Schematic depicting the derivation of the G12 TMZ cell line in vivo. **B)** TMZ IC_50_ curves are derived from treatment response of G12 PL and G12 TMZ cells cultured in 2D. **C)** IC_50_ values reported for the G12 PL and G12 TMZ cells cultured in 2D.

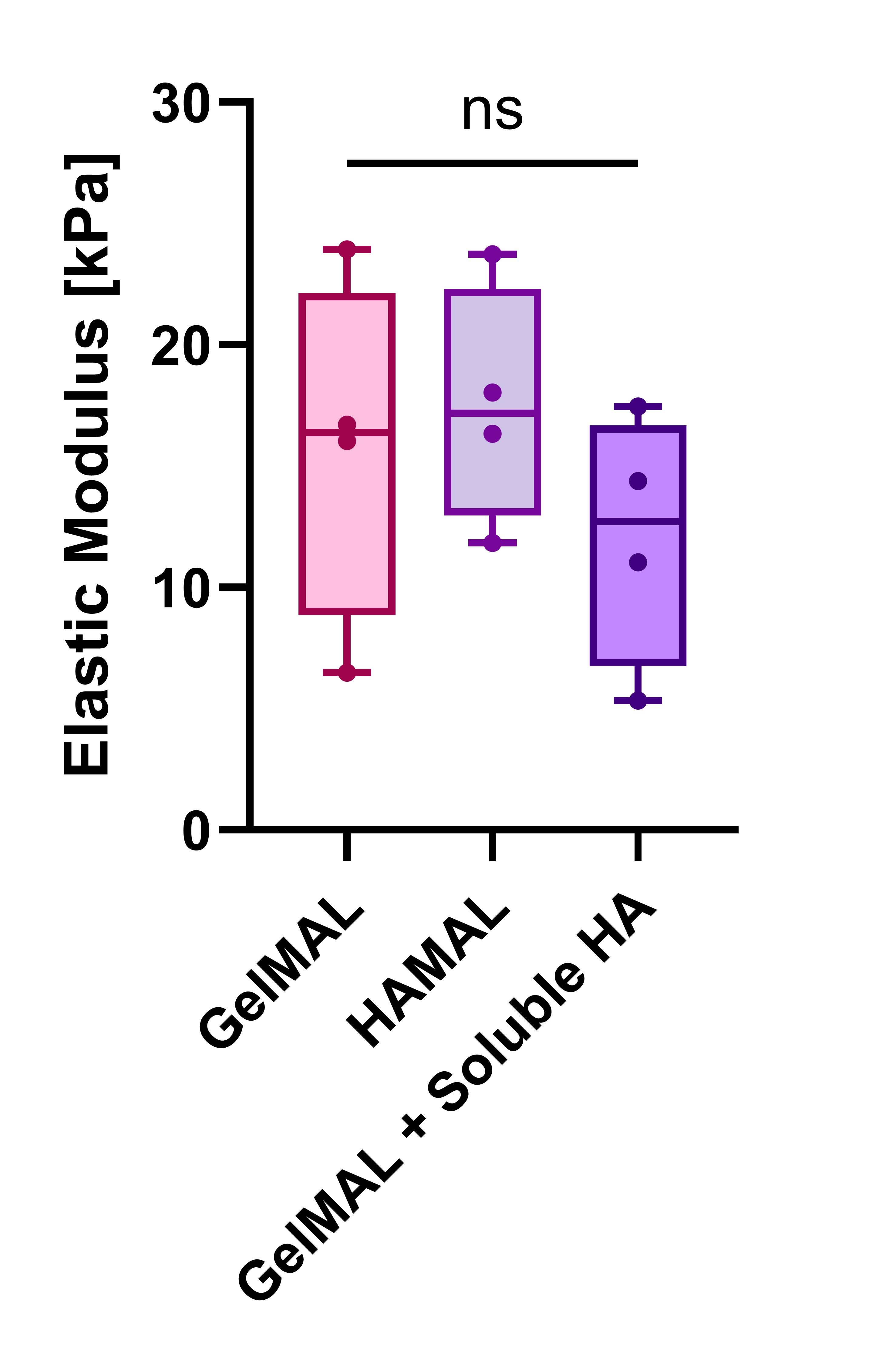

**Figure S3.** Compressive modulus of bulk 4% (w/v) GelMAL hydrogels with or without substituted 0.05% (w/v) HAMAL or cultured in 0.05% soluble HA, a median stiffness of 16.38, 17.19, and 12.71 kPa, respectively. n = 4 hydrogels. Bar plot reports median ± standard deviation. Significance is denoted as *p ≤ 0.05, **p ≤ 0.01, ***p ≤ 0.001, ****p ≤ 0.0001.

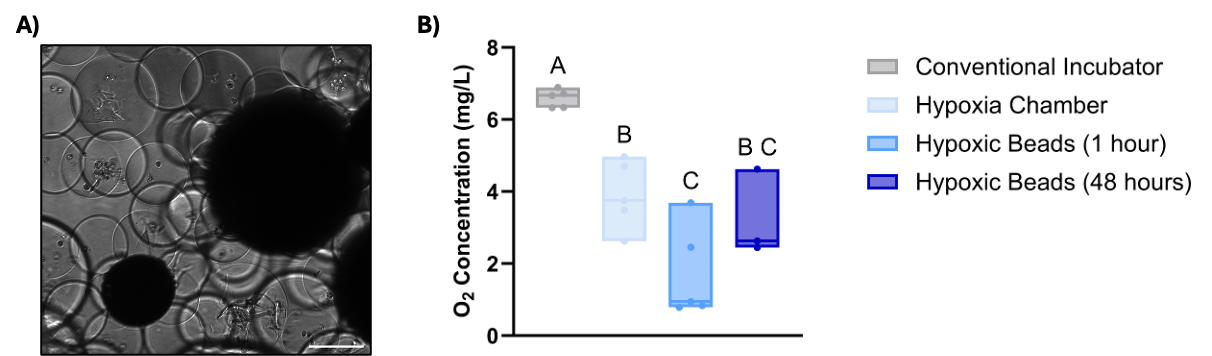

**Figure S4. A)** Brightfield image depicting G12-laden microgels and 0.5% hypoxic beads in culture. **B)** Oxygen concentration in media cultured for a minimum of 2 hours in a traditional incubator (normoxia), 4 hours in a hypoxia chamber (1% O_2_), and 1 hour or 48 hours with hypoxic beads (0.5%) in a traditional incubator. Significance is denoted as *p ≤ 0.05, **p ≤ 0.01, ***p ≤ 0.001, ****p ≤ 0.0001.
